# Characterisation of *Marteilia cocosarum* in the Wash Estuary, UK, linked to mass mortalities of cockles (*Cerastoderma edule*), and its relationship to closely related species

**DOI:** 10.1101/2025.02.13.638039

**Authors:** Anna Tidy, Ron Jessop, Georgia M. Ward, Matthew J. Green, Kelly S. Bateman, David Bass, Jasmine E. Hunt, Stuart H. Ross, Chantelle Hooper

## Abstract

Globally, *Marteilia* parasites have been associated with significant mass mortality events in populations of commercially important bivalve molluscs, frequently resulting in large-scale fishery collapses and substantial socio-economic impacts. The Wash Estuary, UK, supports several bivalve fisheries, and among these, common cockles *Cerastoderma edule* have suffered unusually high mortalities since 2008. We investigate potential causes of these mortalities, and confirm infection with *Marteilia cocosarum*, strongly associated with cockle moribundity, also confirming its presence in archived samples collected in 2009. Molecular and light microscopy screening of samples collected during mortality events in 2021, including healthy (buried) and moribund (weak, unable to bury) cockles, indicated high prevalence of *M. cocosarum* in moribund cockles (PCR incidence up to 95%) in contrast to healthy cockles (up to 42%), suggesting an association between cockle moribundity and *Marteilia* infection. Analysis of the full ribosomal RNA array identified consistently different nucleotides between *M. cocosarum* infections in The Wash (denoted as genotype WE) and those in Wales (denoted genotype BI). 83% of infections in The Wash could be identified as *M. cocosarum* WE and 12% as *M. cocosarum* BI, with both genotypes recovered from 5% of infected animals. Histopathologically, *M. cocosarum* WE infects the gill, mantle and connective tissues, identical to observations of *M. cocosarum* infecting Welsh cockles. Ongoing cockle mortalities in The Wash raise concerns regarding the sustainability of this resource ecologically and economically. Additional measures may be required to reduce the spread of this pathogen, noting that its distribution beyond The Wash and Wales is currently unknown.

## 1. Introduction

*Marteilia* (Ascetosporea, Paramyxida) species are of worldwide significance owing to their detrimental impacts on bivalve mollusc hosts, particularly those of economic importance. Nine species are currently assigned to the genus (Skujina et al., 2022), affecting hosts including oysters (*Ostrea, Crassostrea* and *Saccostrea* spp.), mussels (*Mytilus, Modiulus* and *Xenostrobus*), cockles (*Cerastoderma* spp.) and clams (*Ruditapes, Scrobicularia, Solen* spp.), as most recently reviewed by Ward et al., (2016). The mode of infection and pathogenicity is known to vary depending on the host, with most species infecting the digestive gland (Berthe et al., 2004), and several *Marteilia* species are notorious for their deleterious effect on cultivated bivalve hosts.

In European populations of the common cockle *Cerastoderma edule*, *Marteilia* parasites have previously caused mass mortalities in two Spanish regions, and were linked to mortalities in Wales, United Kingdom. The cockle fisheries of the Ría de Arousa, Galicia, Spain, were previously amongst the most productive shellfisheries in northwestern Spain, but have suffered tremendous collapses since 2012, linked to infection with a *Marteilia* species, initially designated *Marteilia* “type C”, and later formally described as *M. cochillia* (Carrasco et al., 2012, 2013; Villalba et al., 2014). Following recurrent mortalities in several key Welsh cockle populations since 2002 (Murray & Tarrant, 2015; Woolmer, 2013; Burdon et al., 2014), Skujina et al. (2022) associated these mortalities with infection with a novel *Marteilia* species, *M. cocosarum*, a close but phylogenetically and phenotypically distinct sister species of *M. cochillia*. Unlike most other characterised *Marteilia* species, *M. cocosarum* was shown to primarily infect the base of the gill and mantle tissue, with parasite stages distributed through the connective tissues. This unique tissue tropism distinguished infection with *M. cocosarum* from other species, including *M. cochillia* (Skujina et al. 2022).

Investigation of the distribution of *M. cocosarum* in *C. edule* populations around British and Irish coasts using molecular screening only found evidence of its presence around the Welsh coastline (Skujina et al. 2022). *M. cocosarum* was found at high prevalence (48-79%) in cockles across the Welsh sampling sites and was tentatively implicated in cockle mortalities at these locations, in conjunction with additional stressors. However, a lack of data prohibited more conclusive links between parasite infection and mortality events being drawn.

Cockles are known to be infected with many other eukaryotic parasites including metazoans (trematode and nematode worms; copepod and decapod crustaceans), and microbial eukaryotes (including Microsporidia, Alveolata, and other Ascetosporea), as reviewed in de Montaudouin et al., (2021). Despite the large number of eukaryotic parasites known to infect cockles, their prevalence in populations is typically low, and detrimental effects due their presence are unlikely (de Montaudouin et al., 2021). Bacterial and viral agents known to infect cockles are relatively few compared to eukaryotic organisms. The only bacterial species reported to be a threat to cockle production is *Vibrio aestuarianus* (also responsible for mortalities of other bivalve species) which has been identified as the causative agent of cockle summer mortality events in France (Garcia et al., 2021). A retrovirus was initially linked to the presence of disseminated neoplasia (DN) in multiple bivalve species (Romalde et al., 2006), however this has never been confirmed. Despite the causative agents and transmission of DN being currently unknown, DN in cockles has been linked to mass mortalities in Spain (Villalba et al. 2001; Díaz et al. 2016)

The Wash estuary, located on the border of Norfolk and Lincolnshire, United Kingdom, is the largest nature reserve in the UK (Natural England., 2010). Not only is it a hugely important wetland site for wild mammal and bird populations, it also supports numerous molluscan fisheries including *C. edule*, blue mussel *Mytilus edulis* and, to a lesser extent, whelks *Buccinum undatum* (Eastern IFCA., 1992; 2021). Since 2008, cockle stocks in The Wash have suffered unusually high mortalities, particularly affecting larger cockles (>14mm width), leaving beds dominated by smaller juvenile stocks (Eastern IFCA., 2019). These mortalities are considered atypical by the Eastern Inshore Fisheries Conservation Authority (EIFCA) who conduct annual spring surveys and manage the fishery, in that they do not resemble die-offs which routinely occur due to natural events or normal annual variation. Cockle populations have been reported to periodically undergo natural mortality events, including “ridging out” within dense beds, where the larger, less active cockles are forced out of the sediment by younger, more active cockles (Eastern IFCA., 2019). The widespread, atypical losses in Wash cockle populations commonly occur in summer months, during which moribund individuals can be observed gaping on the surface of sandbanks, with slow response to stimuli and overall weakness (Eastern IFCA., 2011). It has also been reported that faster-growing beds appear to be particularly vulnerable to these mortalities (Eastern IFCA., 2011). In fast-growth beds, the cockles reach maturity within 1-2 years, whilst slower-growing cockle beds only produce adults of spawning size after 3-4 years. Atypical die-offs occur in cockles according to size class as opposed to age class, and so those that have reached a marketable size in 2 years on some beds are as vulnerable to mortality as those of 4 years on slower-growing beds. On some fast-growth beds, mortality rates have been observed to exceed 90% of produced biomass once cohorts reach a vulnerable size (Eastern IFCA., 2011). Furthermore, since the events predominantly affect larger size-class cockles, irrespective of age, there has been an overall reduction in the mean cockle size within The Wash, and a shift towards smaller cockles being targeted for harvest (Eastern IFCA., 2022). This could result in the unsustainability of the fishery and reduced longevity of stocks into the future if cockles are harvested before spawning. Moreover, by thinning the stocks at smaller sizes, densities of larger cockles are reduced the following year, perpetuating population reduction. Cockles also hold significant ecological value, particularly through food provision for wild bird populations such as oystercatcher *Haematopus ostralegus* and knot *Calidris canutus* (Stillman et al., 2010), and so widespread population declines also raise the alarm for ecological stability in affected areas.

Cockle mortalities with a similar aetiology have been reported in the Burry Inlet, Wales, UK, since 2004, presenting similar issues for the fishery, and suggesting potential similarities in the causative factor(s) behind these declines (EIFCA, 2011). A comprehensive study of cockle populations in the Burry Inlet was carried out by the Welsh Environment Agency and partners, published by Elliott et al., (2012), which found no direct correlation between the measured water quality parameters and cockle mortality; nor could mortality be conclusively linked to parasite infection as a primary cause, especially as parasite load was found to be higher in populations sampled in the Dee estuary, where mortality incidence was significantly lower. It was concluded that high cockle density, combined with a high energy expenditure and high reproductive output was likely behind population declines, and that more or one other factors (including sediment composition, benthic community composition, water quality or parasite infection) may be contributing to reduced tolerance in cockles already under stress (Elliott et al. 2012). The presence of large foci of haemocyte infiltration was tentatively linked to the presence of haplosporidian parasites (Ascetosporea, Haplosporida), including *Minchinia mercenariae* and *M. tapetis*, though insufficient data were available to confirm this (Longshaw & Malham., 2013). Subsequent studies in Ireland and Spain have confirmed infection of *C. edule* with both species, though infection has not been linked to population declines or mortalities (Albuixech-Martí et al., 2020; Ramilo et al., 2018).

In this study, we 1) Investigate the presence and prevalence of pathogens and pathology potentially associated with cockle mortalities in The Wash by molecular and histopathological screening. 2) Following the detection of *M. cocosarum,* we determine the phylogenetic position of this parasite by sequencing the full ribosomal RNA (rRNA) array. 3) Determine, by revisiting archival cockle specimens from The Wash collected during mortality events in 2009 and 2010, whether *M. cocosarum* was implicated in early cockle mortalities in The Wash, and 4) investigate the presence of other pathogen groups and pathologies previously linked to cockle mortalities, including haplosporidians (*Haplosporidium* and *Minchinia* spp.) and disseminated neoplasia.

## 2. Materials and methods

### 2.1 Sample collection

Samples of approximately 50 moribund *Cerastoderma edule* from the surface of the sandbank and 50 buried “healthy” *C. edule* (not showing clinical disease signs) were collected by the Eastern IFCA from three sites in the Wash Estuary between April and July 2021, and in May 2022. Moribund cockles were identified by weakness, gaping, and the apparent inability to bury in the sediment, while healthy cockles were buried in the sediment and showed normal responses to artificial stimuli (i.e. the two valves of the shell could not be prised apart). Animals were transported overnight in coolboxes to the Cefas Weymouth laboratory for processing. A cross-section comprising representative tissues of all organs was fixed in Davidson’s Seawater Fixative for histopathology and *in situ* hybridisation, and an equivalent section was fixed in RNALater (Sigma Aldrich, USA) for molecular analyses. Additional sections of mantle and digestive gland tissues (approx. 2 mm^3^) were preserved in 2.5% glutaraldehyde in 0.1M sodium cacodylate buffer for transmission electron microscopy (TEM). Archived samples were revisited to determine if the pathogens and pathologies identified in this study were present at the start of the annual mortalities occurring. These samples were collected for histopathology only, as above. All samples included in this study, including sampling date and location, and sample size, are shown in Table 1.

**Table 1.** Summary of samples collected at study sites within The Wash estuary, including site location, sampling date and number of individuals collected. Moribund cockles were collected from the surface sediment, and displayed weakness, gaping and an inability to bury, whereas healthy cockles were dug out of the sediment and showed no clinical signs of reduced health status.

| Site name | Site location | Sampling date | Sample codes | Moribund cockles | Healthy cockles | Total |
| --- | --- | --- | --- | --- | --- | --- |
| Mare Tail | 52° 54' .950" N, 00° 07' .200" E | 28/04/2021 | RA21014 (1-49, 90-139) | 49 | 50 | 99 |
|  |  | 16/05/2022 | MT2022 | 48 | 0 | 48 |
| Dill’s Sand | 52° 56' 150" N, 00° 07' 050" E | 28/04/2021 | RA21014 (50-89, 140-189) | 40 | 50 | 90 |
|  |  | 16/05/2022 | DS2022 | 48 | 0 | 48 |
| Inner West Mark Knock | 52° 50' 300" N, 00° 14' 250" E | 26/07/2021 | RA21022 | 50 | 50 | 100 |
| Bob Halls Sand | 52° 58' 57.6" N 0° 50' 53.88" E | 06/05/2009 | PM19194 | 0 | 13 | 13 |

### 2.2 DNA extraction from RNAlater-fixed tissues

RNALater-fixed cross sections were further dissected to prepare 50 – 100 mg comprising representative tissue from the mantle, digestive gland, gill and adductor muscle. Tissue was homogenised in 800 µl Lifton’s buffer (100 mM EDTA, 25 mM Tris-HCl, 1% SDS, pH 7.5; Winnepenninckx et al (1993)) in a 2 ml Lysing Matrix A homogenisation tube (MP Biomedicals,). Homogenisation was performed in a FastPrep-24 (MP Biomedicals) at 5 m/s for 60 seconds. 20 µl Proteinase K (20 mg/ml; Sigma Aldrich) was added to each homogenate before incubation at 56°C for at least three hours on a static dry block. Following Proteinase K digestion, DNA was extracted from 100 µl of homogenate using the Maxwell RSC Tissue DNA kit, following the manufacturer’s instructions, on a Maxwell 48 Rapid Sample Concentrator (RSC) instrument (both Promega) in batches of 47, with a negative control extraction (i.e. no homogenate added) performed in each batch. Resulting DNA extracts were diluted 1:10 in molecular grade water for use as PCR templates.

### 2.3 DNA extraction from formalin fixed, paraffin-embedded tissues

Paraffin-embedded blocks from samples collected in May 2009 were imaged and measured to calculate the approximate number of 10 µm sections required to obtain 4 mm^3^ tissue. DNA was extracted from sectioned formalin-fixed, paraffin embedded (FFPE) sections using the QIAamp DNA FFPE Advanced UNG kit (Qiagen) using the manufacturer’s protocol including the Uracil-N-Glycosylase (UNG) repair step, with the following modifications: Sections were dissolved in a total volume of 500 µl Deparaffinisation Solution, and incubated at 56°C for 3 minutes to dissolve paraffin was performed twice, in order to fully remove all paraffin from the sections. DNA was eluted into a volume of 50 µl buffer ATE after incubation on the column for 5 minutes, and the eluate reapplied to the spin column and incubation repeated to maximise DNA yield. All optional steps in the manufacturer’s protocol were included, and all “vigorous” vortexing steps were performed at 2700 rpm for 10 seconds. DNA was repaired using the NEBNext FFPE DNA Repair module v2 (New England Biolabs), using a modified version of the manufacturer’s protocol for user-supplied reagents (Section 3). Enzymatic incubation steps were performed as directed, but cleanup steps using sample purification beads were omitted to reduce loss of DNA.

### 2.4 PCR screens and sequencing

The sequences of all primers used in this study are shown in Table 2. All reactions were performed in 20 µl volumes, comprising 1× Green GoTaq Flexi Buffer (Promega), 2.5 mM MgCl_2_ (Promega), 0.4 mM dNTP mix (Meridian Bioscience, USA), 10 µg bovine serum albumin (BSA) (New England Biolabs), 0.5 µM each primer, 0.5 U GoTaq G2 Flexi DNA polymerase (Promega), and 1 µl diluted template DNA. Reaction composition was identical for each round of nested PCR, with 1 µl first round product used as a template for the second round. PCR screens of all *C. edule* DNA extracts were performed using the hemi-nested, paramyxid-targeted 18S ribosomal RNA (rRNA) primer set B (Para1fGW/ParaGenRGW and Para3fGW/ParaGenRGW) and cycling conditions of Ward et al. (2016). A further *Marteilia*-specific nested PCR, targeting the ITS region, was performed to characterise a potentially more phylogenetically informative region of the rRNA array, using the nested primer set (MartDBITSf1/MartDBITSr1 and MartDBITSf2/MartDBITSr2) and cycling conditions of Kerr et al. (2018). Finally, samples collected in 2021 were screened using the nested, haplosporidian-targeted primer set (C5fHap/Sb1n and V5fHapl/Sb2nHap) of Hartikainen et al., (2014), using the modifications presented in Ward et al., (2019). Resultant amplicons were visualised on 1.5% (*w/v*) agarose-TBE gels stained with GreenSafe (NYZ Biotech) alongside 5 µl 100 bp molecular marker (100 bp DNA Ladder; Promega). PCR products were purified using AMPure XP sample purification beads (Beckman Coulter), and bidirectionally Sanger sequenced using the second round primers using the Eurofins Genomics PlateSeq service (Eurofins Genomics).

**Table 2.**
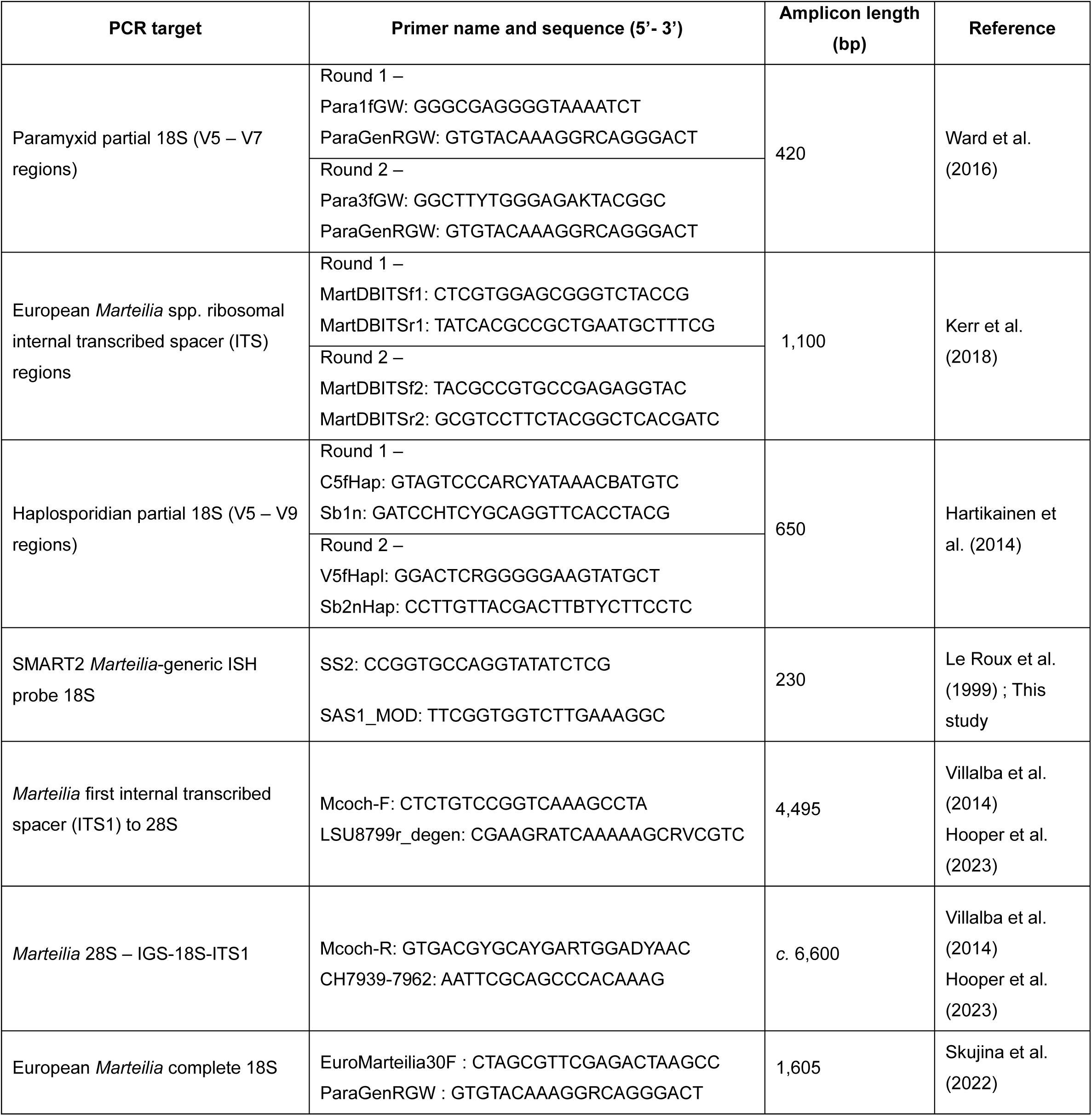
Sequences of primers used in this study for targeted PCR screens and sequencing, ribosomal RNA sequence extension, and generation of digoxigenin-labelled oligonucleotide probes for *in situ* hybridisation.

For paramyxid amplicons that produced mixed chromatogram traces, amplicons were ligated into pGEM-T Easy Vector Systems (Promega) and transformed into *E. coli* JM109 competent cells (Promega) before plated on LB-Ampicillin agar. Transformed colonies were picked and amplified using M13 primers (Messing., 1983), before visualisation, purification and Sanger sequencing as above, using the M13 plasmid primers.

DNA extracts from FFPE material were screened using generic *Marteilia* SSU primers SS2 (Le Roux et al. 1999) and SAS1_MOD, based on the original SAS1 primer of Le Roux et al. (1999) but with an additional C base removed to improve complementarity to European *Marteilia* spp. Reactions were conducted in 20 µl volumes using the same composition as above, but using 5 µl repaired DNA as template. Cycling conditions consisted of an initial denaturation at 95°C for 5 minutes, followed by 45 cycles of 95°C for 30 seconds, 55°C for 1 minute and 72°C for 1 minute. A final extension step at 72°C for 10 minutes was performed, before reactions stored at 4°C. The complete PCR product (i.e. 20 µl) was visualised on 2% (*w/v*) agarose-TBE gels. Resultant bands were excised using a sterile blade and purified using the Monarch DNA Gel Extraction Kit (New England Biolabs). Purified DNA was eluted into a volume of 20 µl, and unidirectionally Sanger sequenced using the forward primer using the Eurofins Genomics TubeSeq Supreme service (Eurofins Genomics, Germany. Resultant chromatogram traces were trimmed to remove primer sequences and low quality bases using MEGA v11 (Tamura et al. 2021), and resultant sequences aligned against full-length small subunit rRNA sequences for European *Marteilia* spp. in order to confirm identity. Due to their short length, these sequences were not included in further analyses.

### 2.5 Marteilia ribosomal RNA array amplification and sequencing

To generate full rRNA arrays for the *Marteilia* lineage detected using group-targeted PCRs, long-range PCR was carried out on PCR-positive individuals (n = 14). Amplicons covering the first internal transcribed spacer region (ITS1), 5.8S gene, second internal transcribed spacer region (ITS2) and partial large subunit 28S gene (hereafter referred to as ITS1-5.8S-ITS2-28S) were amplified using primer Mcoch-F (Villalba et al. 2014) and LSU8799degen (Hooper et al. 2023). Amplicons comprising the 28S gene, intergenic spacer region (IGS), external transcribed spacer (ETS), complete 18S and ITS1 (hereafter 28S-IGS-ETS-18S-ITS1) were amplified using primers Mcoch-R (Villalba et al. 2014) and CH7939-7962 (Hooper et al. 2023). Cycling conditions for both amplicons were as described in Hooper et al. (2023).

Both ITS1-5.8S-ITS2-28S and 28S-IGS-18S-ITS2 amplicons were prepared for Illumina sequencing using the Nextera XT DNA Library Preparation Kit (Illumina) following the manufacturer’s protocol but using half reaction volumes. Libraries were sequenced on a MiSeq Nano reagent kit (500 cycles) on an Illumina MiSeq platform (Illumina). 28S-IGS-18S-ITS1 amplicons were prepared for Nanopore sequencing using the PCR Barcoding Kit (Oxford Nanopore Technologies, UK) following the kit protocol, and sequenced on Flongle flow cells on a MinION Mk1C platform (both Oxford Nanopore Technologies).

Based on the analysis of the full rRNA array sequences, a further PCR covering a region of the array in the 18S gene which discriminated between *Marteilia* identified in The Wash from other *Marteilia* species was performed to demonstrate that differences seen in the full array were consistent across all available samples. The full 18S sequence was amplified using primers EuroMarteilia30F and ParaGenGW, and the cycling conditions presented in Skujina et al. (2022).

### 2.6 Analysis of ribosomal RNA array sequences

Bidirectional Sanger sequence traces generated from 18S paramyxid-targeted and ITS1 *Marteilia*-targeted specific PCRs were trimmed to remove primer sequences and low quality basecalls using MEGA v11 (Tamura et al., 2021), before reads were concatenated and then aligned against existing sequence data for *M. cocosarum* and *M. cochillia* using MAFFT v7.0 (L-ins-i algorithm) (Katoh & Standley., 2013). The resulting alignment was viewed in AliView v1.26 (Larsson., 2014). Indels and base changes in the newly-generated sequences comparative to the two closely-related *Marteilia* species were observed and counted by eye.

Analysis of long-range amplicon data generated using Illumina and Nanopore platforms followed that used by Hooper et al. (2023) for the analysis of *Marteilia* and *Paramarteilia* sequence data.

### 2.7 Phylogenetic analysis

A Bayesian consensus tree was constructed based on all available paramyxid 18S sequence diversity to determine the phylogenetic placement of the *Marteilia* species detected in cockles in The Wash, relative to other paramyxid lineages. A second Bayesian consensus tree, using full transcribed rRNA alignments (ETS-18S-ITS1-5.8S-ITS2-28S) generated in this study (alongside publicly available rRNA arrays for *M. cocosarum, M. cochillia and M. octospora*) was then constructed. Both phylogenies were constructed using MrBayes v3.2.7 (Ronquist et al., 2012) on the CIPRES Science Gateway (Miller et al., 2010). Analysis used two separate MC3 runs, carried out for two million generations with one cold and three hot chains. The first 500,000 generation were discarded as burn-in, and trees sampled every 1000 generations. Resultant consensus tree files were viewed using FigTree v1.4.4 (Drummond & Rambaut., 2007).

### 2.8 Histopathology and Transmission Electron Microscopy (TEM)

Sections of cockle tissue including the adductor muscle, gill and digestive gland were fixed in Davidson’s Seawater Fixative for 24 hours, before transfer to 70% industrial denatured alcohol (IDA) (Pioneer Research Chemicals Limited). Fixed sections were then processed as in Ward et al. (2016), before examination under light microscopy for the presence of parasites and pathologies using a Nikon Eclipse N*i*-E (Nikon). Digital images and measurements were obtained using NIS-Elements imaging software (Nikon).

Post-fixation of glutaraldehyde-preserved digestive gland and mantle tissues was carried out in 1% osmium tetroxide/0.1M sodium cacodylate buffer before processing as in Ward et al. (2016). Grids were examined using a JEOL JEM1400 transmission electron microscope (JEOL) and digital images captured using an AMT XR80 camera with AMT V602 software (AMT Imaging).

### 2.9 In situ hybridisation (ISH)

Digoxigenin (DIG)-labelled oligonucleotide probes targeting the 282 bp “SMART2” region of the *Marteilia* 18S gene were synthesised from genomic DNA extracted from a *M. cocosarum*-infected cockle using primers SS2 (Le Roux et al., 1999) and SAS1_MOD. Probes were synthesised in 100 µl PCR reactions comprising 1× Green GoTaq Flexi Buffer, 2.5 mM MgCl_2,_ 0.2 µM each primer, 200 µM PCR DIG labelling mix (Roche, Switzerland), 2.5 U GoTaq Flexi DNA polymerase, and 6 µl template DNA. Cycling conditions were as detailed in Le Roux et al. (1999) for the SS2/SAS1 primer set. Resultant amplicons were visualised as above on 2% agarose-TBE gels before probes were purified by the addition of 1× reaction volume of AMPure XP sample purification beads, washed twice with 80% molecular-grade ethanol, and eluted into 50 µl molecular-grade water. Probe was quantified using QuantiFlour ONE dsDNA reagents on a Quantus fluorometer (both Promega) and normalised to a concentration of 10 ng/µl.

Whole animal sections of *Marteilia*-infected cockles collected in 2021, 2022 and 2009 (3 µm thickness) were collected onto poly-L-lysine coated glass slides. Sections were dewaxed and rehydrated with Xylene substitute (Thermo Fisher Scientific) twice for five minutes each, followed by two washes in 100% ethanol for 4 minutes each. Sections were then washed once in Tris-buffered saline (TBS) at pH 7.5 for 2 minutes. Next, sections were overlaid with 1 ml Proteinase K in TBS pH 7.5 (45 µg/ml) for 20 minutes at 37°C, before washing in deionised water for 5 minutes. Sections were then treated with ice-cold 12% acetic acid for 45 seconds to increase permeability of the tissue, before dehydration by washing in 70% ethanol for 3 minutes, followed by 100% ethanol for a further 3 minutes. Sections were rinsed in 2× saline sodium citrate (SSC) for 1 minute with agitation, before storage in TBS (pH 7.5). GeneFrames (Thermo Fisher Scientific) were applied to each slide, and sections overlaid with 120 µl hybridisation buffer (50% formamide, 4× SSC, 1× Denhardt’s solution, 10% dextran sulphate, 250 µg/ml single-stranded DNA from salmon testes) containing 50 ng probe. For negative control samples, no probe was added to the hybridisation buffer. Sections were denatured by heating to 94°C for 6 minutes, before immediately cooling on ice for a further 6 minutes. Slides were then hybridised overnight at 42°C in a humid chamber. The following morning, GeneFrames were removed and sections washed with 3 times for 10 minutes each with wash buffer (0.5× SSC, 6 M urea, 20 µg/ml BSA) pre-warmed to 42°C, before two further washes for 5 minutes each in TBS pH 7.5. Sections were then overlaid with a milk blocking buffer (1× TBS pH 7.5, 6% (*w/v*) non-fat milk powder (Sigma Aldrich)) for 1 hour at room temperature. Following this, sections were washed twice for 5 minutes each in TBS pH 7.5, before incubation overlaid with Anti-DIG-AP Fab fragments (Roche) diluted 1:300 in TBS pH 7.5 for 1 hour in darkness. To remove excess antibody, slides were washed 5 times for 10 minutes each in TBS (pH 7.5), before transfer to TBS pH 9.5 for two minutes. Slides were then overlaid with 5-bromo-4-chloro-3-indoyl-1-phosphate (BCIP) nitroblue tetrazolium (NBT) chromogenic substrate (Roche), diluted 20 µl/ml in TBS pH 9.5, for up to 1 hour in darkness at room temperature. After this time, sections were washed three times in TBS pH 9.5 for 3 minutes each, followed by counterstaining with Bismarck Brown Y solution (5% in 35% ethanol) for 15 minutes. Sections were then gently rinsed in running tap water to remove excess counterstain, and dehydrated by washes in 70% ethanol and 100% ethanol (30 seconds each), followed by two rinses in xylene substitute for 1 minute each. Sections were then air-dried for a minimum of 30 minutes, before cover slipping. Sections were then examined under light microscopy using a Nikon Eclipse NI (Nikon, Japan) and images of labelled parasite cells in host tissue captured using NIS-Elements Imaging Software (Nikon, Japan).

## 3. Results

### 3.1 PCR prevalence of parasites in Wash cockles

All Wash cockle samples (n=385) were initially screened for paramyxids using a nested, 18S-targeted primer set, producing 198 amplicons (51%). The result of these PCR screens, including the number of PCR positive animals per site and per health status (i.e. moribund or healthy) is summarised in Table 3, and detailed below. Phylogenetic analysis of the resulting sequences found that 22 of these were 100% identical to 18S sequences for *M. cocosarum* from the Burry Inlet, Wales. We denote this sequence type “*Marteilia cocosarum* BI1” (Burry Inlet 1). 167 samples amplified a closely related but distinct *Marteilia* sequence type with no identical matches in the NCBI GenBank database, most closely related to *M. cocosarum* (sequence identity 99.4%). We denote this sequence type “*Marteilia cocosarum* WE1” (Wash Estuary 1). Nine samples produced mixed chromatogram traces which required cloning, and subsequent sequencing of cloned amplicons revealed both *M. cocosarum* BI1 and *M. cocosarum* WE1 to be amplified from these animals.

**Table 3.**
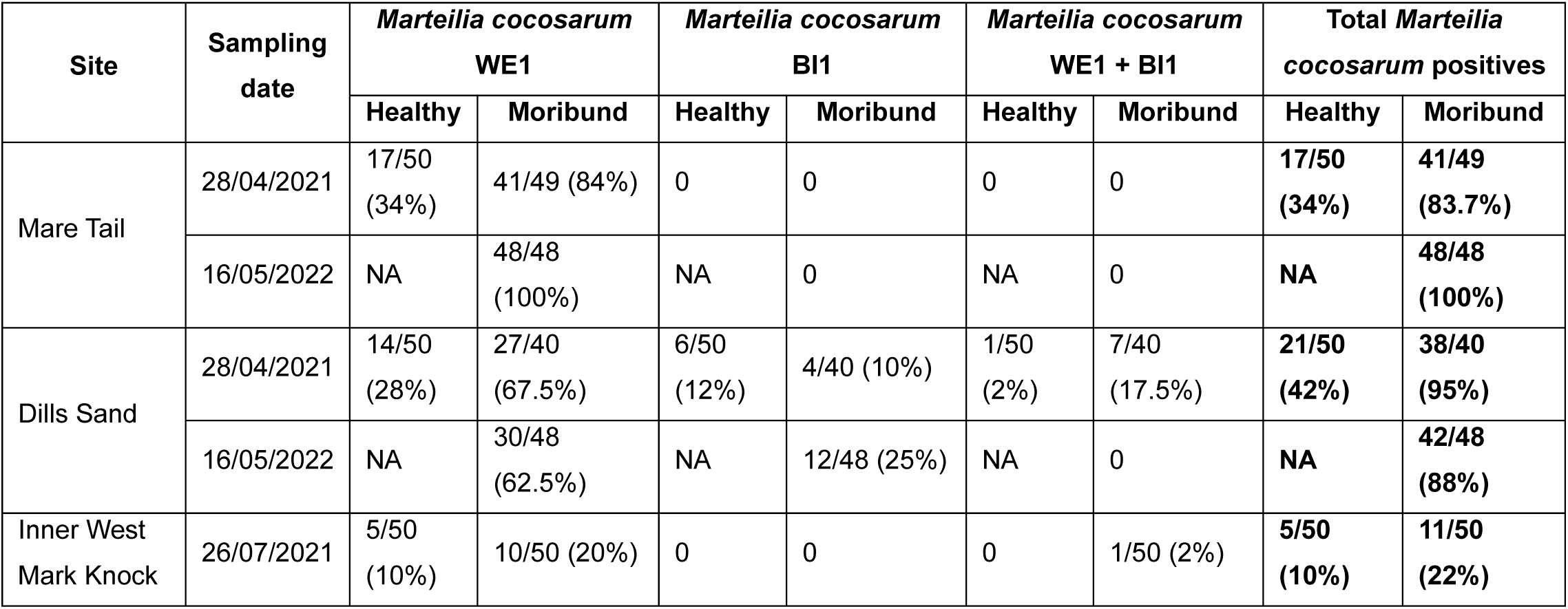
Total *Marteilia cocosarum* prevalence in healthy vs. moribund cockles per site, detected via PCR screen.

The highest proportion of *Marteilia*-positive animals were collected from the Mare Tail and Dills Sand sites, in samples collected in 2021 and 2022. A greater number of moribund animals were PCR positive than healthy animals at both Mare Tail (83.7% vs 34% in 2021) and Dills Sand (95% vs 42% in 2021). No healthy cockles were collected in 2022, but 100% of moribund animals from Dills Sand and 88% of moribund animals from Mare Tail were PCR positive. At Inner West Mark Knock, sampled in July 2021, 22% of moribund animals were PCR-positive, compared to 10% of healthy animals. Across all sites, there were significantly greater *Marteilia* positives (Chi-Sq p = 0.00001158) detected by PCR in moribund cockles compared to healthy cockles.

PCR screens of cockle extracts samples collected in 2021 using a nested primer set targeting the 18S gene of haplosporidians produced a total of 158 amplicons (54%), with no significant difference in the total number of positives between healthy or moribund animals at any site. A greater number of haplosporidian positives were found from the sites sampled in April (Mare Tail (total 55/98); Dills Sand (total 66/90)) than Inner West Mark Knock, sampled in July (17/100). Sequencing of these amplicons produced a total of 128 usable sequences, with the remining 18 amplicons producing unusable, mixed chromatogram traces, likely indicating the amplification of more than one haplosporidian sequence type. In total seven haplosporidian sequence types were amplified, as summarised in Table 4. The most abundant haplosporidian lineage amplified across all three sites were *Minchinia* spp., including *M. tapetis*, a parasite of the grooved carpet shell *Ruditapes decussatus* also known to infect *Cerastoderma edule* in Spain (Carballal et al., 2020), *M. tapetis* was present at all sites, and in both healthy and moribund animals. Prevalence was highest at Dills Sand, with a similar percentage of positives between moribund (27.5%) and healthy (22%) animals. An uncharacterised *Minchinia* species, previously shown to have a strong association with *C. edule* populations in Spain (Ramilo et al., 2018), was also present at all sites, generally at a higher PCR prevalence in moribund animals. *Minchinia* sp. was amplified from 57%, 55% and 12% of moribund animals from Mare Tail, Dills Sand and Inner West Mark Knock, respectively. The parasite was also present in healthy animals from each site, at 26%, 42% and 4% prevalence, respectively.

**Table 4.** Total Haplosporidian prevalence in healthy vs. moribund cockles per site, detected via PCR screen.

| Site | Sampling Date | Health status (total animals) | *Minchinia tapetis* | *Minchinia* sp. ex. *Cerastoderma edule* | *Minchinia chitonis* | *Haplosporidium edule* | Coastal water clone KF208577 (Fleet Lagoon) | Coastal water clone MT334478 (Mumbles Pier) | Novel sequence type 01 |
| --- | --- | --- | --- | --- | --- | --- | --- | --- | --- |
| Mare Tail | 28/04/2021 | Moribund (49) | 2 (4%) | 28 (57%) | 0 | 0 | 0 | 0 | 0 |
|  |  | Healthy (50) | 5 (10%) | 13 (26%) | 2 (4%) | 2 (4%) | 1 (2%) | 1 (2%) | 1 (2%) |
| Dill's Sand | 28/04/2021 | Moribund (40) | 11 (27.5%) | 22 (55%) | 0 | 0 | 0 | 0 | 0 |
|  |  | Healthy (50) | 11 (22%) | 21 (42%) | 0 | 1 (2%) | 0 | 0 | 0 |
| Inner West Mark Knock | 26/07/2021 | Moribund (50) | 5 (10%) | 6 (12%) | 0 | 1 (2%) | 0 | 0 | 0 |
|  |  | Healthy (50) | 3 (6%) | 2 (4%) | 0 | 0 | 0 | 0 | 0 |

Five other haplosporidian sequence types were also amplified from cockles, though at much lower prevalence. The *C. edule* parasite *Haplosporidium edule* was amplified from a low number of healthy cockles from Mare Tail and Dills Sand (4% and 2%, respectively), and a single moribund cockle (2%) from Inner West Mark Knock. The chiton parasite *Minchinia chitonis* amplified from two healthy cockles collected at Mare Tail (4% prevalence), but no other site. A previously detected but uncharacterised sequence type, originally amplified from water and sediment collected in the Fleet Lagoon, Dorset, UK, was amplified from a single healthy cockle from Mare Tail. A second environmental sequence type, originally detected in water samples collected close to the Mumbles Pier, Swansea, UK, was also amplified from a single healthy cockle at Mare Tail. A novel sequence type was amplified from a single cockle, also a healthy individual collected at Mare Tail. This sequence has no close characterised relatives, though showed high identity (99%) to an uncharacterised lineage previously amplified from incubation water associated with the crab species *Carcinus maenas* and *Cancer pagurus* collected in Newton’s Cove, Dorset, UK.

### 3.2 Histopathology, in situ hybridisation and Transmission Electron Microscopy (TEM) of Marteilia cocosarum

*Marteilia*-like cells were observed within the connective tissues, most commonly observed in the gill and mantle tissues of 40% of the cockles sampled from Mare Tail and 37% of cockles sampled from Dills Sand. *Marteilia*-like cells presented with the typical cell within cell arrangement associated with paramyxid infections and were often associated with large focal areas of haemocytic infiltration (Fig 1). In the case of systemic infections *Marteilia*-like cells were observed within the haemal sinuses and the vesicular connective tissues surrounding the digestive gland and gonad but were not observed within the epithelial cells of the digestive gland tubules and stomach tissues. ISH labelling confirmed presence of *Marteilia*-like cells within areas of pronounced haemocytic infiltration, especially in the connective tissues of the gills and mantle, parasite cells staining a deep blue colour (Fig 2).

**Figure 1.**
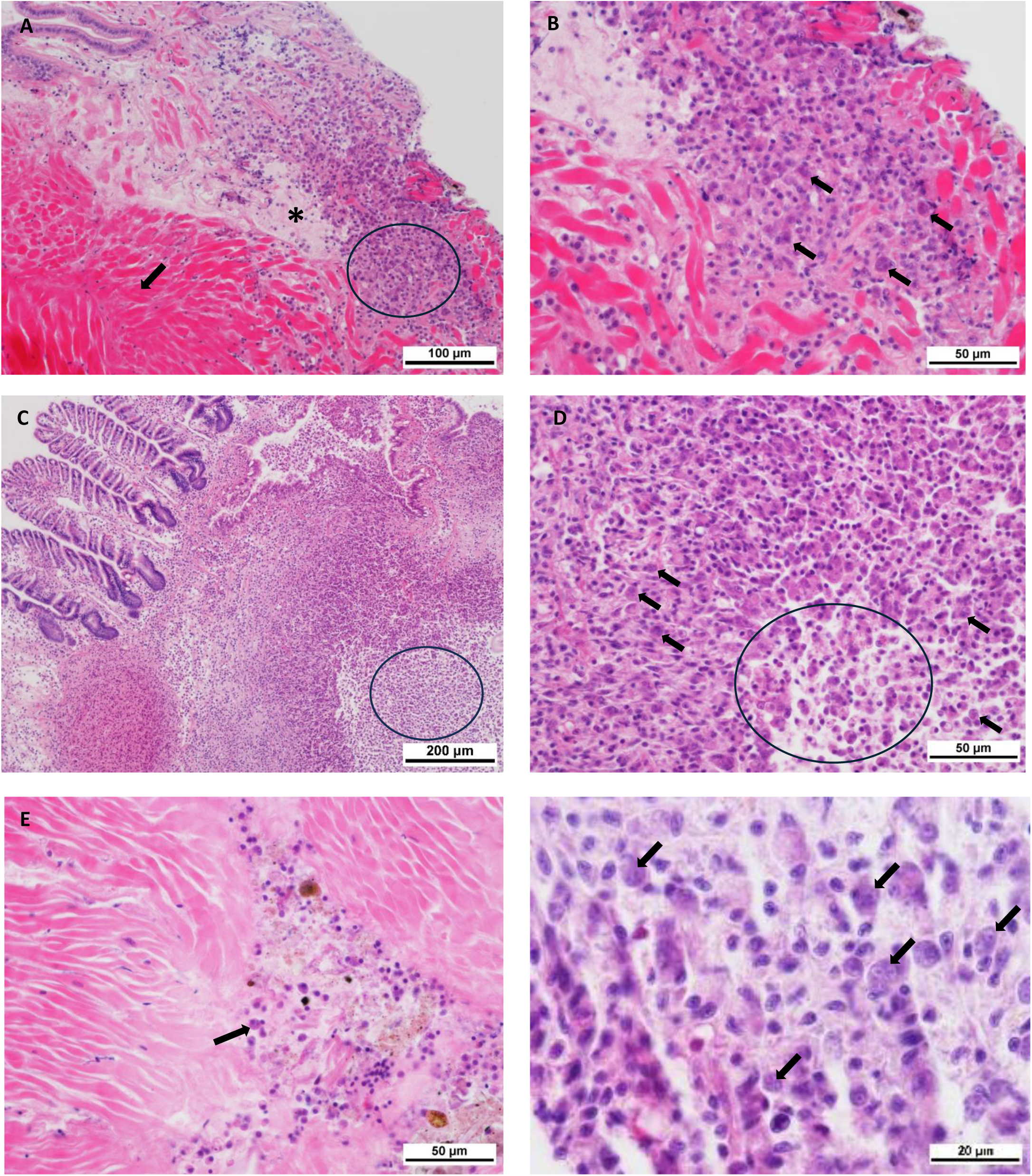
A) Muscle (arrow) and vesicular connective tissues (*) of *C. edule* displaying a focal area of haemocytic infiltration (circle). Scale bar = 100 µm. B) Higher magnification of ‘A’, focal area of haemocytic infiltration containing large multinucleate cells (arrows), cell within cell arrangement can be observed which is typical of paramyxid infections such as *Marteilia* sp. Scale bar = 50 µm. C) Marked host response observed in the Gill area with dense haemocyte infiltration containing multinucleate *Marteilia*-like cells, tissue necrosis (circled area) was also observed. Scale bar = 200 µm. D) Higher magnification of ‘C’ cellular inflammation, large multicellular *Marteilia*-like cells (arrows) are visible throughout the lesion. Area of necrosis (circle) shows sloughing, rounding and dissociating cells associated with apoptosis and necrosis. Scale bar = 50 µm. E) *Marteilia*-like cells (arrow) observed in the sinus of the adductor muscle. Scale bar = 50 µm. F) Cell within cell appearance of *Marteilia*-like cells (arrows) in connective tissues of gill. Scale bar = 20 µm. All images H&E stained.

**Figure 2.**
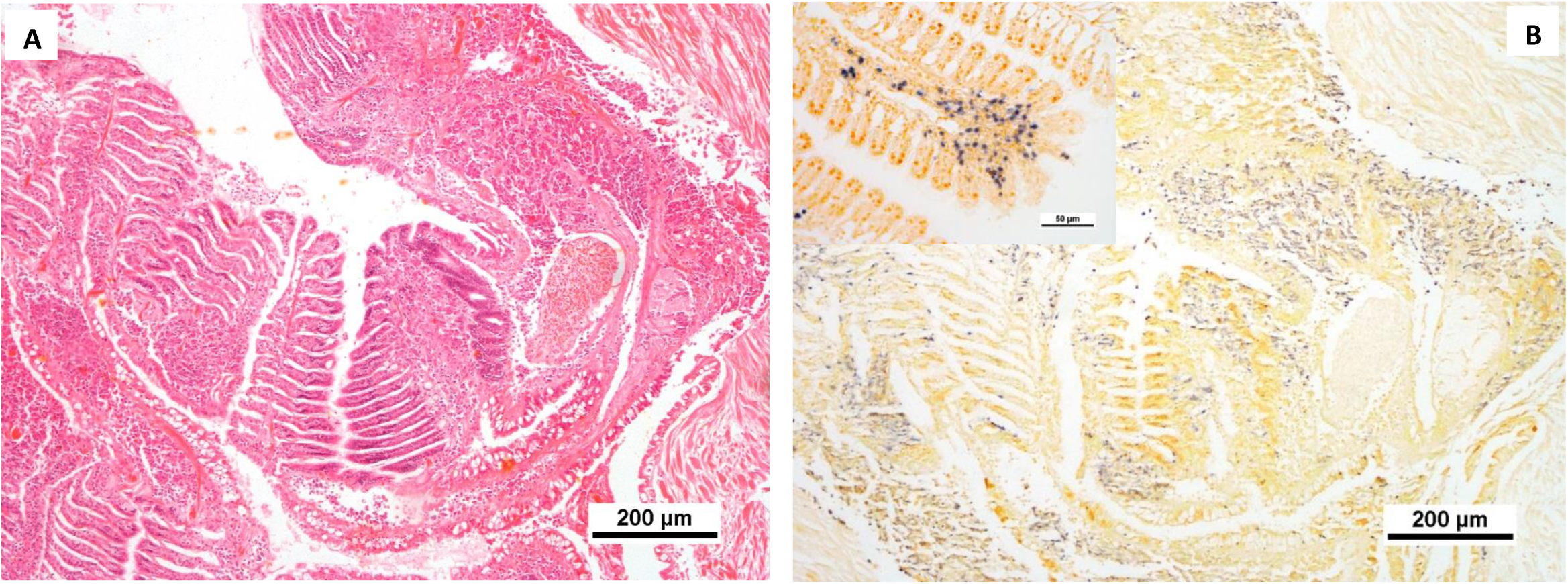
*In situ* hybridisation (ISH) of *Marteilia*-like parasites within *C. edule* tissues. A) Negative control gill tissues. H&E stain. B) *In situ* hybridisation (ISH) of the same region of gill tissues, labelled with Digoxigenin (DIG)-labelled Smart-2-probe targeting the 282 bp region of the Marteilia 18S gene. Blue stain highlights presence of *Marteilia*-like parasites within the connective tissues and associated with pronounced areas of haemocytic infiltration. Scale bars = 200 µm. Inset displaying magnified region of gill tissue infected by *Marteilia*-like pathogens. Scale bar = 50µm.

As described by Skujina *et al*. (2022) primary cells appeared to contain up to three tertiary cells, no spore like formations were noted. TEM analysis found multiple *Marteilia*-like cells distributed throughout the connective tissues, clearly displaying the characteristic cell within cell arrangement of the secondary and tertiary cells within the primary cell (Fig. 3). Haplosporosome-like structures were observed within primary and secondary cells only. Distinct double membranes were observed separating the primary, secondary and tertiary cells, primary cell membrane denser in appearance when compared to secondary and tertiary cells. Interestingly the cell membrane of the primary cell appeared to possess pore like structures, small discrete areas where the membrane appeared as a single membrane, it is not clear whether these areas may be involved in endocytosis between the host cells and parasite. Higher levels of infection were observed in moribund animals when compared to healthy animals at both sites, 60% and 63% prevalence in moribund animals and 24% and 12% in healthy animals at Mare Tail and Dills Sand respectively.

**Figure 3.**
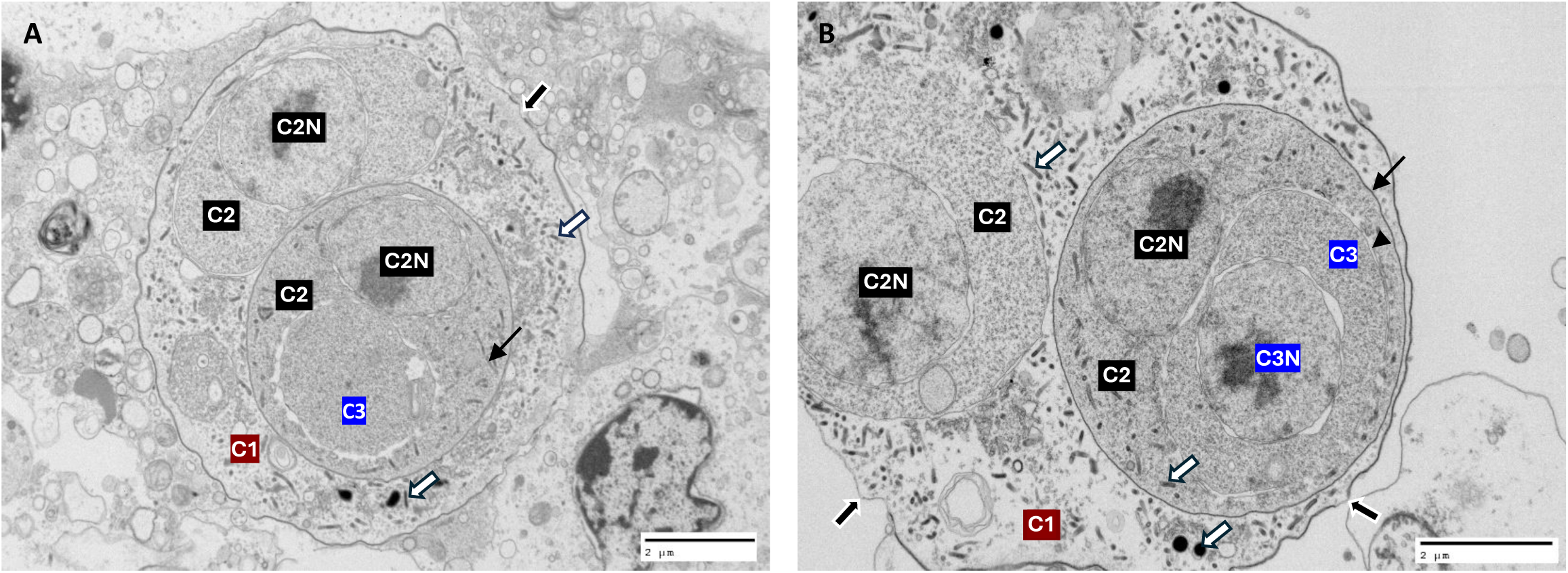
Transmission electron microscopy of *Marteilia*-like cells within *C. edule* tissues. A) Single primary cell (C1) containing multiple rod shaped haplosporosomes (white arrow) and two secondary cells (C2) containing clear nuclei (C2N). Some secondary cells contained haplosporosomes (line arrow) and tertiary cells (C3). Note the pores within the primary cell membrane (black arrow). B) Higher magnification detailing secondary and tertiary cells. Primary cell (C1) contains two secondary cells (C2) both of which display clear nuclei (C2N). Secondary cells are bound within a clear membrane (line arrow) and can contain haplosporosomes (white arrows). Tertiary cell (C3) also contains a clear nucleus (C3N) and is bound within a clear membrane (arrowhead). Haplosporosomes were not observed within the tertiary cells. Note the appearance of single membrane in certain areas of the primary cell membrane (black arrows). Scale bars = 2 µm.

Cockles at both sites showed multiple co-infections with various different pathogens by histology, summarised see Table 5 for summary of histology results. *Nematopsis* sp. is a common infection of cockles and was observed at both sites during this study, although higher level prevalence was reported at Mare Tail. Rickettsia-like organisms were observed within the digestive gland at both sites at similar prevalence and a bacterial infection was noted in some cockles. This appeared to be seen more frequently in moribund animals and was associated with necrosis of the tissues so is likely a secondary infection. Necrosis and inflammation was observed in a large number of animals samples from Mare Tail when compared to Dills Sand. Inflammation was commonly associated with presence of Marteilia-like cells. Haemocytic neoplasia was observed in a few cockles at each site, the majority thought to be type B (Carballal et al., 2001; Iglesias., 2006; Metzger et al., 2006). *Bucephalus minimus* was observed at low level at both sites in healthy and a few moribund animals at Dills Sand. *Trichodina* sp. were also observed at both sites in healthy animals and a few moribund animals at Mare Tail. Low levels of Turbellaria sp., digenean and ciliate species were observed at both sites.

**Table 5.** Summary of pathogens and pathology observed in healthy and moribund cockles, and total cockles examined histologically at each site.

| Site and condition | *Marteilia*-like cells | *Nematopsis* sp. | RLO/CLO | *Trichodina* sp. | *Bucephalus* sp. | *Turbellaria* sp. | Digenean | Ciliates | Bacteria | Haemocytic neoplasia | Inflammation | Necrosis |
| --- | --- | --- | --- | --- | --- | --- | --- | --- | --- | --- | --- | --- |
| **Mare Tail** |  |  |  |  |  |  |  |  |  |  |  |  |
| Moribund cockles (49) | 60% | 88% | 15% | 22% | 0 | 10% | 0 | 1% | 25% | 3% | 78% | 28% |
| Healthy cockles (50) | 24% | 60% | 14% | 14% | 16% | 2% | 2% | 0 | 0 | 0 | 24% | 0 |
| Total cockles (99) | 40% | 72% | 14% | 18% | 9% | 6% | 1% | 0 | 11% | 1% | 37% | 12% |
| **Dills Sand** |  |  |  |  |  |  |  |  |  |  |  |  |
| Moribund cockles (40) | 63% | 24% | 6% | 0 | 2% | 0 | 0 | 0 | 4% | 0 | 0 | 0 |
| Healthy cockles (50) | 12% | 66% | 16% | 12% | 8% | 6% | 4% | 4% | 0 | 6% | 42% | 0 |
| Total cockles (90) | 37% | 45% | 11% | 6% | 5% | 3% | 0 | 2% | 2% | 3% | 21% | 0 |

### 3.3 Molecular analyses and phylogeny of Marteilia in the Wash

The Wash *Marteilia* lineage (*M. cocosarum* WE1) can be phylogenetically distinguished from the closely-related genotype *M. cocosarum* BI1 and more distantly related sister species *M. cochillia* using a relatively short (approx. 500 bp) region of the 18S gene of the rRNA array, using Sanger sequencing. In this region, *M. cocosarum* WE1 shows five consistent nucleotide differences to *M. cocosarum* BI1, and four in comparison to *M. cochillia* (Table 6). To further discriminate *M. cocosarum* WE1, additional PCRs and high-throughput sequencing were undertaken to retrieve sequence data for the full transcribed rRNA array. Across the full array, the two *M. cocosarum* lineages show 99.83% similarity, with 14 consistent nucleotide differences (Table 6). Though the ITS regions (ITS1 and ITS2) typically show higher rates of variation between closely related lineages, comparison of these regions showed there to be only one nucleotide difference between *M. cocosarum* WE1 and *M. cocosarum* BI1 in ITS1 (681 bp), and five differences within the ITS2 (781 bp), the same number of consistent differences observed within the 18S gene.

**Table 6:** Nucleotide differences and percentage similarity across the different gene regions of the ribosomal rRNA array between *Marteilia* lineages. Sizes of each region of the array are given.

| Lineages compared | Nucleotide differences/percentage similarity |  |  |  |  |  | % similarity over full transcribed array |
| --- | --- | --- | --- | --- | --- | --- | --- |
|  | ETS<br>1,547 bp | 18S<br>1,792 bp | ITS1<br>681 bp | ITS2<br>781 bp | 28S<br>3,471 bp | Full transcribed array<br>8,414 bp |  |
| *M. cochillia* ~<br>*M. cocosarum* WE1 | 13<br>(99.16%) | 3<br>(99.83%) | 6<br>(99.12%) | 6<br>(99.23%) | 9<br>(99.74%) | 37 | 99.56 |
| *M. cochillia* ~<br>*M. cocosarum* BI1 | 15<br>(99.03%) | 4<br>(99.78%) | 6<br>(99.12%) | 9<br>(99.85%) | 10<br>(99.71%) | 44 | 99.48 |
| *M. cocosarum* BI1 ~<br>*M. cocosarum* WE1 | 2<br>(99.87%) | 5<br>(99.72%) | 0<br>(100%) | 5<br>(99.36%) | 1<br>(99.97%) | 13 | 99.85 |

Comparatively, *M. cocosarum* WE1 was 99.56% similar to *M. cochillia* from Spanish cockles, with 37 nucleotide differences across the full array. The greatest sequence divergence between these two lineages is within the ITS1 with six nucleotide differences (99.12% sequence identity) and ETS with 13 nucleotide differences (99.16% sequence identity). Between the two *M. cocosarum* lineages, only two nucleotide differences exist within the ETS region, and one within the ITS1 region. Given this level of sequence similarity, the 18S gene or ITS2 regions appear to be most suitable for discriminating between the three lineages (Table 6), although an inconsistent indel in the ITS2 of *M. cocosarum* (Skujina et al. 2022) makes this region difficult to use as a diagnostic marker, due to the requirement to perform confirmatory sequencing. Therefore, we suggest the 18S gene as the most appropriate option for discrimination between lineages. Full transcribed rRNA arrays were deposited to GenBank under accession numbers PP853058-PP853071.

Due to 100% sequence identity across all 18S sequences generated in this study for *M. cocosarum* WE1, a single representative sequence was used for phylogenetic analysis. Bayesian phylogenetic analysis shows that *M. cocosarum* WE1 and BI1 branch as sister clades with very high posterior probability support (0.99) (Fig. 4). The *M. cocosarum* WE1 + BI1 clade branches as sister to *M. cochillia*, though this relationship receives lower statistical support (posterior probability 0.90). In phylogenetic analyses of the full ribosomal arrays of *M. cocosarum* WE1 (n=13), BI1 (n=6) and *M. cochillia* (n=6), using *M. octospora* as an outgroup, each lineage forms a mutually exclusive, monophyletic clade with maximal statistical support (posterior probability = 1.0) (Fig. 5). *M. cocosarum* WE1 and *M. cocosarum* BI1 group as sister clades (with maximal support), clearly distinguishing the two lineages as independent genotypes, with *M. cocosarum* lineages in turn grouping distinctly from *M. cochillia* from Spanish cockles.

**Figure 4.**
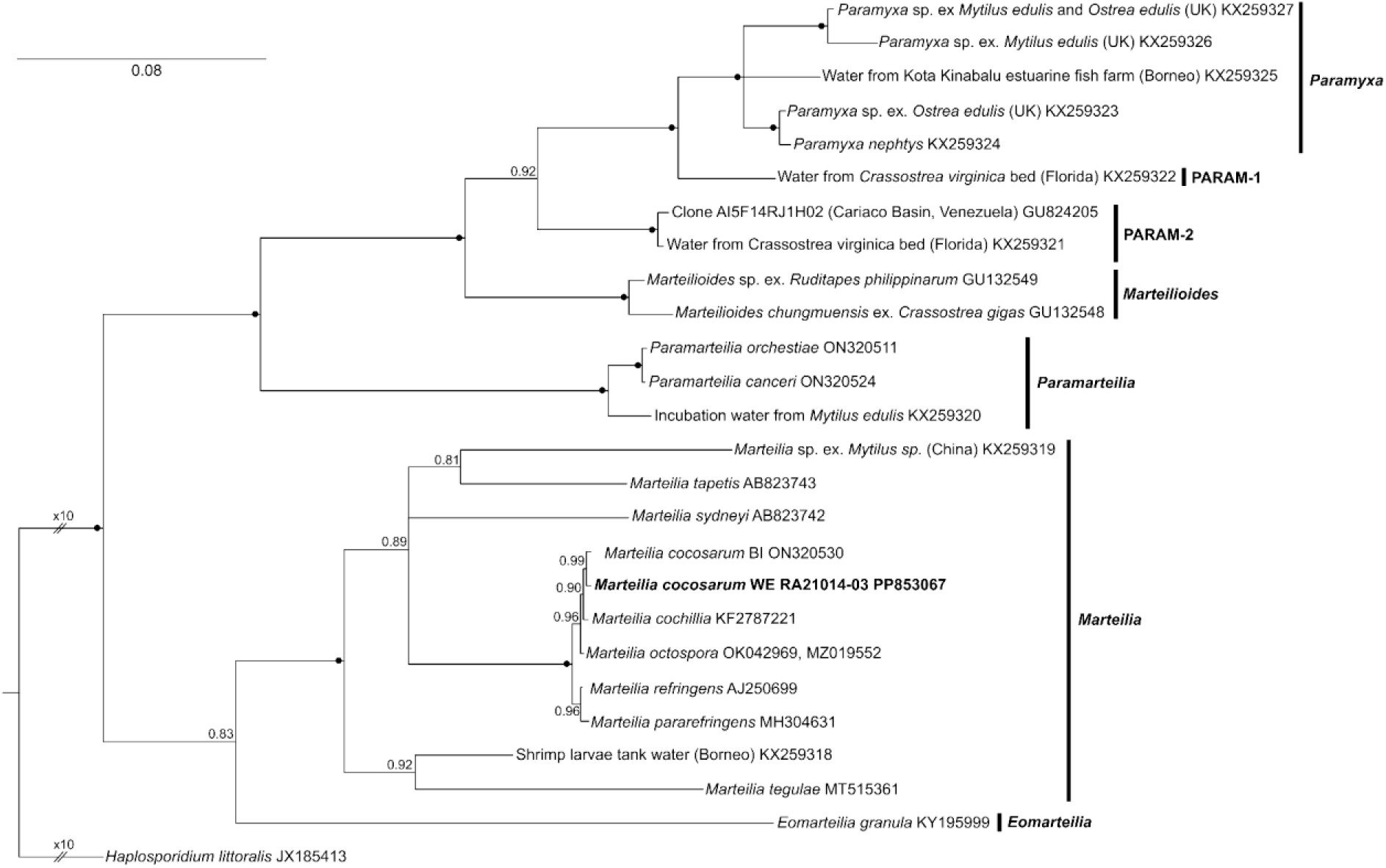
A phylogenetic comparison of closely related Paramyxid lineages including *M. cocosarum* WE1 (in bold) and *M. cocosarum* BI1 both infecting *C. edule* in the UK, as well as *M. cochillia* known to infect *C. edule* in Spain. Phylogenetic tree constructed using Bayesian analysis using 18S region sequences of the rRNA array and *Haplosporidium littoralis* as an outgroup. Maximal Bayesian Posterior Probabilities support values are labelled above branches. Accession numbers are detailed following the species/sample name or ID.

**Figure. 5.**
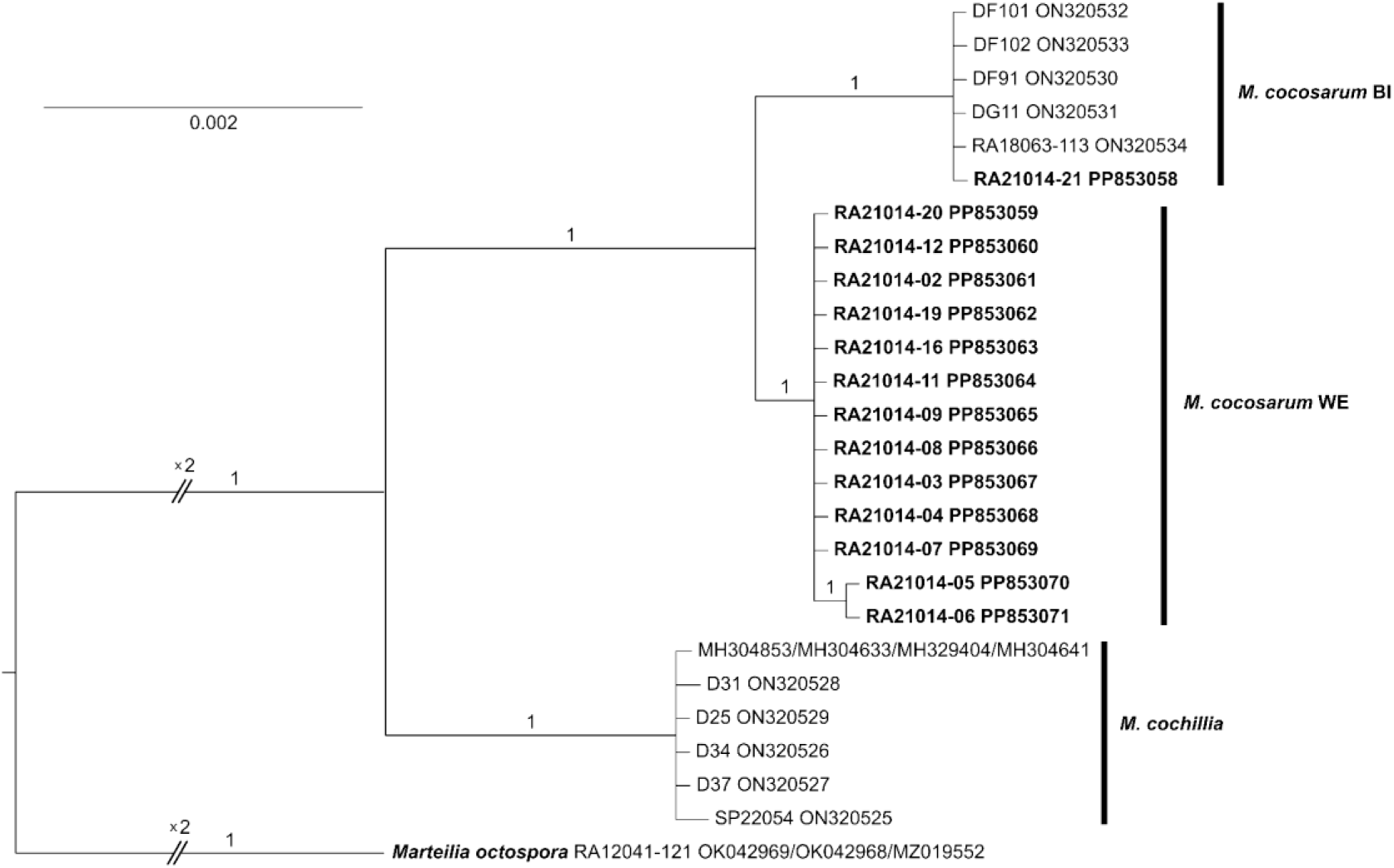
A phylogenetic comparison of *Marteilia cocosarum* WE infecting *C. edule* in the Wash estuary, to *Marteilia cocosarum* BI infecting *C. edule* in the Bury Inlet, and their relationship to *Marteilia cochillia* infecting *C. edule* in Spain. Based upon Bayesian analysis of rRNA sequences with *Marteilia octospora* as an outgroup.

## 4. Discussion

The present study provides evidence of a novel genotype of the paramyxid parasite *Marteilia cocosarum* infecting common cockles *Cerastoderma edule* in The Wash Estuary, UK, which we denote *M. cocosarum* WE1 (Wash Estuary 1). Prevalence of the parasite, using both molecular and histopathological screening techniques, was observed to be significantly higher in moribund, surface-bound cockles than healthy, buried animals. Reanalysis of archive samples collected at sites within The Wash experiencing mortalities in 2009 using microscopy and molecular techniques shows the parasite to have been present within cockle populations within the estuary since at least May 2009, shortly after mortalities were first documented in 2008.

Molecular and histopathological analyses confirm this lineage to show high identity to *M. cocosarum* BI1 (Burry Inlet 1), from the Burry Inlet, Wales, both genetically and in its pathology and mode of infection. *M. cocosarum* genotypes are demonstrably distinct phylogenetically and histopathologically from their closest known relative *M. cochillia*, which infects *C. edule* in Spain. Analysis of full rRNA array sequences allows the two *M. cocosarum* genotypes to be distinguished by consistent nucleotide differences in several regions, with the two lineages showing high sequence identity to each other across the rRNA array (99.83%), and lower similarity to *M. cochillia* (99.56%). The ETS within the intergenic spacer region showed the greatest sequence divergence between *M. cocosarum* and *M. cochillia*, but low variation was observed between the two *M. cocosarum* genotypes (15 nucleotide differences compared to two over the 1,547 bp region). To distinguish between *M. cocosarum* WE1 and *M. cocosarum* BI1, we recommend sequences of the 18S gene, which shows five consistent nucleotide differences between the two *M. cocosarum* genotypes, and three to four nucleotide differences between *M. cochillia and M. cocosarum,* dependent on genotype (Table 6). Although ITS2 shows a larger number of consistently variable nucleotides, an inconsistent indel in this region makes it difficult to retrieve consistent quality sequences without the need for amplicon cloning (Skujina et al 2022).

Histopathological examination showed that *M. cocosarum* WE1 infections are consistent with those observed for *M. cocosarum* BI1. Parasite cells are observed throughout the gill and mantle tissues, and within the connective tissues of the adductor muscle and digestive gland (but not within the digestive gland tubules themselves) in heavily infected animals. Both genotypes also elicit a host response, with inflammatory regions containing haemocytic infiltration observed in infected individuals. This tissue tropism differs from that observed with other *Marteilia* species, including *M. cochillia*, which are typically found within the epithelium of the digestive gland tubules, between digestive epithelial cells of the digestive ducts, and within the intestinal lumen (Carrasco et al. 2015; Villalba et al., 2014).

The initial characterisation of *M. cocosarum* followed widespread mortalities of *C. edule* in the Burry Inlet, Wales, and within populations along the Welsh coast. Although the parasite was detected at high PCR prevalence, and high levels of infection in individual animals was observed by histopathology, across the sites surveyed, only a tentative link between parasite presence and mortality could be made, with the parasite implicated alongside a number of additional stressors (Skujina et al. 2022). These conclusions were made primarily due to a lack of mortality data for many sites, and incomplete sampling limiting histopathological and electron microscope examination of affected cockles. Furthermore, cockles collected during the Welsh study were not recorded as either moribund or “healthy” at the time of sampling, and therefore the correlation between *M. cocosarum* presence and cockle health status could not be explored. In our present sample set, a strong association between cockle moribundity and *Marteilia* infection can be made: weak, gaping cockles on the sediment surface show significantly higher infection prevalence and intensity than those buried within the sediment and showing no clinical disease signs.

*Marteilia* prevalence in moribund cockles was lower at Inner West Mark Knock (IWMK) compared to the other two sites surveyed (22% vs 95% and 83.7% for Dills Sand and Mare Tail in 2021, respectively). Prevalence was always higher in moribund relative to healthy cockles. The extent of mortalities on IWMK in 2021 did not appear to be dissimilar to that of Dills Sand or Mare Tail, with regards to losses in cockle numbers and biomass. Equally, IWMK was sampled later on in the summer season than the other two sites, when temperatures were likely to be warmer and therefore *Marteilia* prevalence would be greater due to increased stress. Yet, the lack of *Marteilia* prevalence here strongly suggests that additional agents are involved in the cockle mortalities observed. It is important to consider the influence of other stressors on cockle populations, particularly across hydrographically distinct areas of The Wash estuary. Exposure to varying biotic (e.g. cohabiting benthic invertebrates, other parasites) and abiotic (e.g. chemical exposure or nutrient limitation) conditions may reduce overall host fitness, resulting in increased susceptibility to parasite infection. The involvement of planktonic crustaceans as intermediate hosts has long been considered to play a role in the life cycle of *Marteilia* species (Audemard et al., 2002; Arzul et al., 2014), and so consideration may be given to the composition of benthic and planktonic invertebrate communities across The Wash.

Histopathology identified disseminated neoplasia in all three populations surveyed. Prevalence was highest at Inner West Mark Knock, with the condition observed in 16 individuals, 15 of which were moribund animals, and one was a healthy animal. Disseminated neoplasia is a leukaemia-like cancer reported in at least 15 species of marine bivalves, including clams, mussels, oysters and cockles (Metzger et al., 2016; Diaz et al., 2011).

Typically, the disease is characterised by the presence of large, anaplastic cells within the haemolymph. Two different types of neoplastic cells were observed within the Wash cockles, described as Type A and B, characterised by the shape of nuclei and arrangement of cells and distinguished phylogenetically as two distinct cancer lineages (Carballal et al., 2001; Metzger et al., 2016). The majority presented signs of Type B neoplasia (12 individuals), whilst 4 individuals were observed to have Type A. In diseased animals, the cells multiply and reduce the number of normal haemocytes, eventually resulting in the death of the host (Barber., 2004). It is therefore possible that the higher incidence of disseminated neoplasia observed at Inner West Mark Knock, where *Marteilia* prevalence was comparatively low compared to the other two sites surveyed, may be linked to the atypical mortalities observed at this site. Disseminated neoplasia in cockles has also been identified in at least one site in Wales (Bruzos et al., 2023), but its link to the mortalities occurring in this region, to our knowledge, has not been investigated.

Cockles were also screened by PCR for the presence of haplosporidian parasites, after previous studies suggested a link between a number of haplosporidian species and cockle declines in the Burry Inlet (Elliott et al. 2012; Longshaw & Malham 2013). Two lineages – *Minchinia tapetis* and an uncharacterised *Minchinia* sp. closely related to *M. mercenariae* – were amplified, with both showing higher prevalence in moribund than apparently healthy cockles. However histopathological screening detected only a single haplosporidian infection at low intensity in a healthy animal from Mare Tail (Table 5). This same animal was PCR-positive for *M. tapetis*. Both *Minchinia* lineages amplified in this study have been previously reported to be present in cockle populations in Europe. *M. tapetis* has been observed infecting *C. edule* in Galicia, Spain, where infection prevalence appeared high, but intensity within infected individuals was low (Carballal et al. 2020). Seasonal studies in Spain showed infection prevalence to be linked to seawater temperature, with infection highest in the warmer months. In our study, molecular prevalence of the parasite was highest in animals sampled from Dills Sand in April, however as no repeat sampling was conducted at the same sites in the summer it cannot be determined whether the prevalence of observable (histopathological) infection is higher among cockles in The Wash estuary during the warmer months. The second *Minchinia* lineage amplified in this study is also known to occur within *C. edule* populations within Galicia. *Minchinia* sp. appears in cockles at a lower prevalence, and infections were of moderate or high intensity, with haplosporidian cells observed throughout the connective tissues in the digestive gland, gills and gonad. A strong host response is observed, though to date is not associated abnormal cockle mortalities. Uni- and bi-nucleate parasite stages and multinucleate plasmodia were observed in infected individuals, but no spore stages could be seen, preventing conclusive identification of the parasite (Ramilo et al. 2018). However, given the high sequence similarity of this lineage to *M. mercenariae*, it is likely that this is the same parasite previously reported associated with cockle populations experiencing abnormal mortalities and population declines, including those in the Burry Inlet (Elliott et al. 2012; Longshaw & Malham, 2013). As no link between haplosporidian infection and cockle moribundity could be made at any site surveyed in this study, and that none of the haplosporidian lineages found in The Wash have been demonstrably linked to increased or unusual cockle mortalities, it is unlikely that these parasites are a primary cause of mortalities within Wash cockles.

## 5. Conclusion

The combined analyses from the present study describe a novel genotype of *M. cocosarum* WE1 infecting cockles in The Wash estuary. This discovery follows the recent description of a closely related genotype of the same parasite *M. cocosarum* BI1 causing marteiliosis in UK cockles in the Burry Inlet, Wales (Skujina et al., 2022). We also provide evidence that *M. cocosarum* has been present in cockle populations within The Wash since at least May 2009, which is shortly after documented evidence of mortalities and population declines were reported in 2008. Further work is required to understand the geographic extent of the parasite within The Wash and surrounding areas, including cockle fisheries in the northeast and the Thames catchment. Also important is understanding the risk of transmission of the parasite to new locations, particularly through intentional or accidental translocation of bivalve species between production areas. Further work should also aim to understand any differences in pathogenicity between this novel genotype compared to *M. cocosarum* BI1, as well as the biotic and abiotic factors which may contribute to high parasite prevalence and increased likelihood of infection outbreak.

## 6. Acknowledgements

This study was funded by Defra project AHPFFB2: Diagnostics and Emerging Disease.

## Notes

### Competing Interest Statement

The authors have declared no competing interest.

### Summary of Updates

Revised to include the below reference in the reference list. Previously accidentally omitted from the paper. Kerr, R., Ward, G.M., Stentiford, G.D., Alfjorden, A., Mortensen, S., Bignell, J.P., Feist, S.W., Villalba, A., Carballal, M.J., Cao, A. and Arzul, I., 2018. Marteilia refringens and Marteilia pararefringens sp. nov. are distinct parasites of bivalves and have different European distributions. Parasitology, 145(11), pp.1483-1492.

